# Warming-induced switches in dominance are built into intraguild predation systems

**DOI:** 10.64898/2026.06.18.733167

**Authors:** Peter Kamal, Emanuel A. Fronhofer

**Affiliations:** ISEM, University of Montpellier, CNRS, IRD, EPHE, 34095 Montpellier, France

## Abstract

Warming affects food webs globally. In the iconic intraguild predation food web module consisting of a basal resource, a specialist consumer, and an omnivorous predator, resource enrichment can favor the predator by increasing the relative importance of intraguild predation compared to resource competition. Here, we integrate empirically established thermal scaling relationships into a model of intraguild predation. We show that warming can shift the power balance between consumer and predator and affect invasion and equilibrium outcomes by inducing changes to resource enrichment – without any differences in thermal optima between species. The nature of these shifts depends on the thermal scaling of resource self-regulation and the strength of resource top-down regulation. We also test the capacity of several generic early warning signals to predict these shifts and find variance-based indicators to be more reliable than autocorrelation-based ones. Our results have implications for predictive food web ecology and biocontrol applications under global change.

## Introduction

Climate change affects global biodiversity through rises in both mean and variance of temperatures (Bellard et al., 2012; Parmesan and Yohe, 2003; Vázquez et al., 2015). Changing temperatures can cause species to shift or expand their range and invade habitat that was previously too cold (Chen et al., 2011; Sentis et al., 2021), often leading to large biodiversity losses (Kamal et al., 2025; Thompson and Fronhofer, 2019). Warming can also shift the dynamics of local species interaction networks (Binzer et al., 2012; Bideault et al., 2019), a phenomenon often referred to as “rewiring” (Blanchard, 2015). In food webs - trophic interaction networks -, previous work has found many different mechanisms for such rewiring, including thermal asymmetries between species (Gibert et al., 2022), behavioral responses of mobile generalists (Bartley et al., 2019), interactions with concurrent nutrient enrichment (Bonnaffé et al., 2024), and changes in available phenotypic variation (Barbour and Gibert, 2021). Here, we focus on a classic food web module to show that such rewiring can be built into the way a simple system responds to warming.

Intraguild predation (IGP) is a ubiquitous trophic interaction (Polis et al., 1989; Arim and Marquet, 2004; Thompson et al., 2007) and classically comprises three species: a basal resource, a specialist consumer feeding on the resource, and an omnivorous predator feeding on both the resource and the consumer (Holt and Polis, 1997). This simple configuration exhibits a remarkable richness of dynamics due to the interplay of competitive and predatory interactions (Polis and Holt, 1992). IGP systems can also stabilize larger food webs that they are a part of, by adding interior attractors and making chaotic dynamics less likely (McCann and Hastings, 1997; Borrvall et al., 2000). Therefore, this three-species module has become one of the fundamental building blocks of our mechanistic understanding of food webs. It also has an important application: Natural enemies that engage in IGP are often used as biocontrol agents against herbivore pests (Polis and Holt, 1992; Martin et al., 2013). Understanding IGP systems is therefore of great interest for both conceptual and applied issues.

The majority of IGP studies has focused on the local dynamics, and in particular on the effects of resource enrichment (Morin, 1999; Diehl and Feißel, 2000; Mylius et al., 2001; Amarasekare, 2008), which causes a switch in system behavior: At low resource levels, intraguild competition for resources is the main driver of the system and leads to exclusion of the weaker competitor. When resources are enriched, intraguild predation becomes the main driver of the system and the omnivore excludes the specialist, while there is a coexistence region at intermediate resource levels. These basic dynamics can be mediated by other factors, such as spatial dynamics (Amarasekare, 2006) or adaptive foraging (Křivan and Diehl, 2005).

How IGP systems respond to temperature has so far only received limited attention. Temperature is a key driver of almost all biological processes (Kingsolver, 2009) and plays a prominent role in the Metabolic Theory of Ecology. This theory posits that the metabolic rate governs biological processes at all levels from cells to ecosystems (Brown et al., 2004). The metabolic rate is highly susceptible to changes in temperature (Gillooly et al., 2001), and thus, higher-level processes such as movement (Gibert et al., 2016) and digestion (Sentis et al., 2013) are impacted by changes in temperature as well. Accordingly, warming has been shown to impact virtually all rates that determine the dynamics of food webs (Rall et al., 2012; Gilbert et al., 2014; Uiterwaal and DeLong, 2020). Despite this wealth of general knowledge, we could identify only four studies exploring the effect of temperature on IGP systems specifically. Three empirical papers (Sentis et al., 2014; Frances and McCauley, 2018; Rogers et al., 2018) all documented short-term increases in consumption rates as temperature increased, including increases in intraguild predation rates. They conclude that the system should switch from competitive to predatory dynamics as temperature increases, potentially leading to destabilization of the food web or consumer exclusion. The fourth paper, a modelling study (Thunell et al., 2021), shows the opposite effect: By including age-structure and ontogenetic diet shifts in the predator, they show that warming should in most cases lead to predator exclusion.

Despite these first advances, a comprehensive understanding of the effects of temperature on IGP systems is still lacking. All of the empirical work described above (Sentis et al., 2014; Frances and McCauley, 2018; Rogers et al., 2018) used short-term feeding trials to estimate consumption rates. Forecasts based on such estimates can be biased because they neglect the impact of other, unmeasured parameters (Briggs and Borer, 2005). It is especially surprising that the temperature dependence of the resource’s dynamics has only been peripherally explored (Thunell et al., 2021), despite the generally recognized importance of resource enrichment for IGP systems (Morin, 1999; Diehl and Feißel, 2000). Interplay between the different trophic levels could impact the response of the entire system to warming, as processes on the resource level differ in their temperature sensitivity from processes on higher trophic levels: The consumption and death rates of consumers tend to be less sensitive to temperature changes than the rates of resource population growth and self-regulation (Brown et al., 2004; Rall et al., 2012; Davis et al., 2026). These scaling asymmetries could produce a switch in system behavior across temperatures if they have the potential to change the relative enrichment of the resource, and, in turn, the relative importance of competitive versus predatory intraguild interactions (see Fig. 1A for an illustration). Consequently, warming could cause a shift from consumer dominance to predator dominance or vice versa – without the additional (and trivial) mechanism of a difference in thermal optima between the two species.

**Figure 1:**
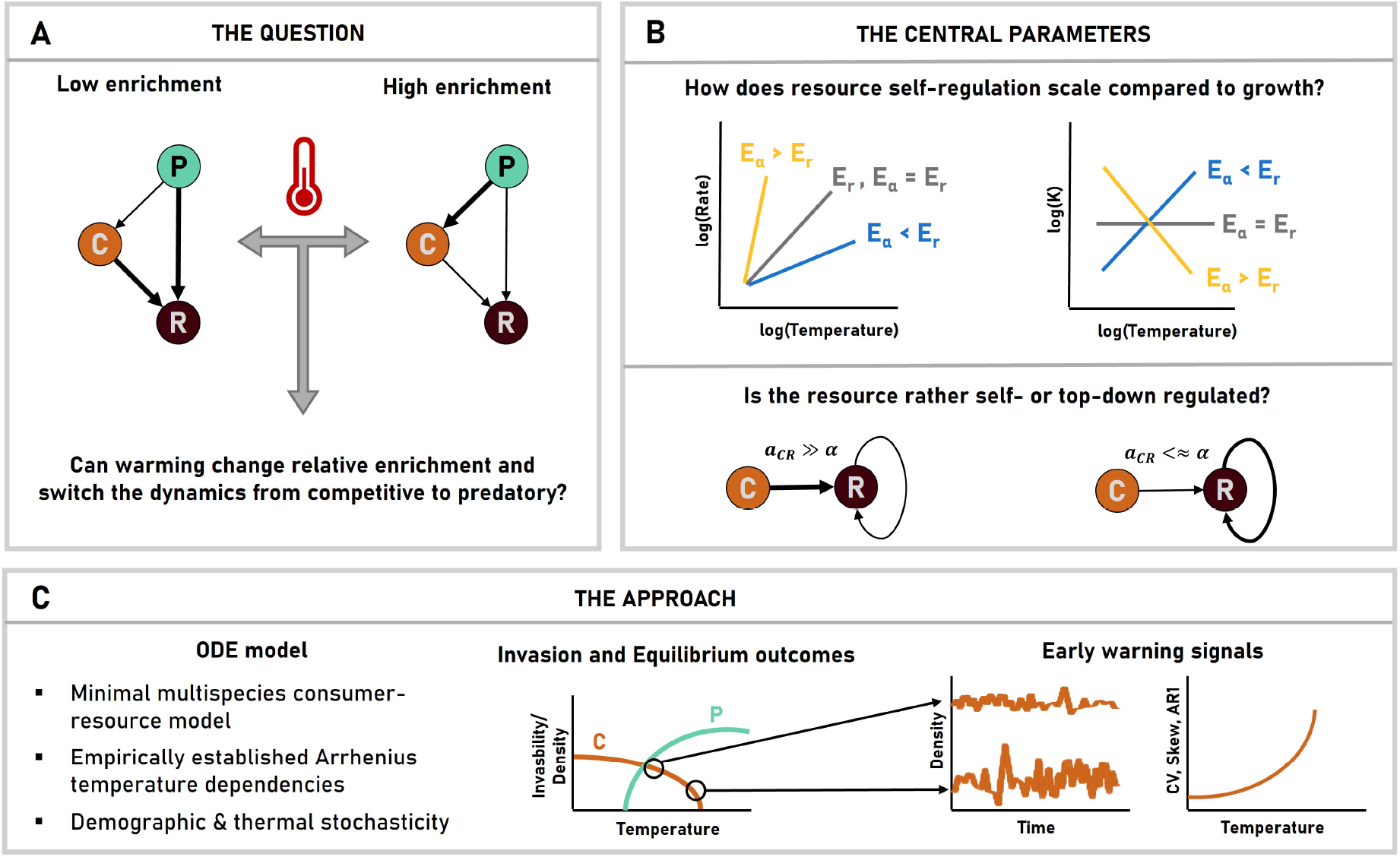
Conceptual illustration of the research question, the central parameters, and the methodological approach. (A) A classical IGP system consisting of a basal resource *R*, a specialist consumer *C*, and an omnivorous predator *P* . The thickness of the arrows indicates the dominant intraguild process under different resource levels (Diehl and Feißel, 2000; Mylius et al., 2001). (B) Central parameters under study. The upper panel shows different scenarios for the qualitative temperature scaling of the resource self-regulation rate *α* compared to its maximum growth rate *r* and the consequences for the resource’s resulting single-species equilibrium density *K*; all of the illustrations are for the rising parts of the thermal performance curves. The lower panel contrasts systems in which the resource is rather top-down regulated (left) with systems in which the resource is rather self-regulated (right) by varying the thickness of arrows. (C) Qualitative illustration of potential changes in invasibility or equilibrium density of consumer or predator across temperatures (left), expected behavior of the consumer’s population time series at two different temperatures (middle), and expected behavior of the corresponding early warning signals across temperatures (right). Closer to the thermal extinction/non-invasibility threshold, the time series should exhibit more variance, first-order autocorrelation, and skew (Scheffer et al., 2009; Dakos et al., 2012a).

Here, we explore whether such temperature-driven switches in the power dynamics of IGP systems occur, and if so, under which conditions. We integrate empirically established temperature dependencies of essential biological rates into a minimal model of IGP. We analyze the model using invasion analysis as well as deterministic and stochastic simulations. We find that the hypothesized switches can indeed occur, and that their nature depends crucially on two parameters: (i) the thermal scaling of resource self-regulation, and (ii) the relative strength of resource top-down regulation. Because of the large amount of empirical data required to quantitatively predict these switches, we subsequently test whether generic early warning signals (Scheffer et al., 2009; Dakos et al., 2012a) can reliably predict them. Fig. 1 summarizes the question, our key parameters, and our methodological approach.

## Model description

We model a local food web comprised of a resource *R*, a specialist consumer *C*, and an omnivorous predator *P* . The system dynamics at temperature *T* are described by the following set of ordinary differential equations:

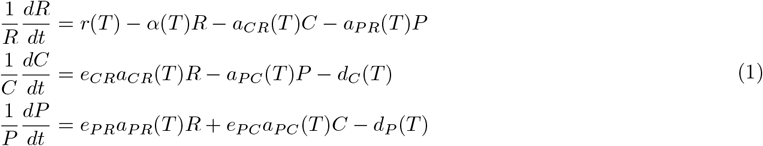

The resource grows logistically with maximum population growth rate *r* and self-regulation coefficient *α* (Mallet, 2012). The consumer feeds on the resource with a consumption rate *a*_*CR*_, and the predator feeds on both the resource and the consumer with consumption rates *a*_*P R*_ and *a*_*P C*_, respectively. Consumers and predators convert their resources into offspring with a species-pair-specific conversion efficiency *e*_*ij*_, and die with a species-specific death rate *d*_*i*_. This is a three-species extension of a classic consumer-resource model with a linear functional response (Diehl and Feißel, 2000). Throughout our analysis, we assume that the consumer is the stronger competitor for the resource; otherwise, the predator always excludes the consumer (Holt and Polis, 1997).

### Temperature dependencies

We make each parameter *θ* of the system temperature-dependent following an Arrhenius equation (Davis et al., 2026):

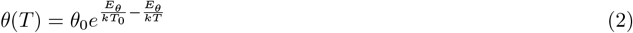

where *θ*_0_ is the rate at the baseline temperature *T*_0_ = 20^*°*^*C* (293.15K), 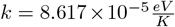 is the Boltzmann constant, and *E*_*θ*_ is the rate-specific activation energy that determines the slope of the scaling relationship. As such, our model is designed to capture the dynamics of well-mixed communities of ectotherms across a temperature range in which all rates are in the rising part of their thermal performance curves and there are no differences in thermal optima between the species. We do not model the falling parts of the thermal performance curves as there are few established models and estimates for deactivation energies of specific rates (Dell et al., 2011).

We use the most recent estimates of activation energies for ectotherms (Davis et al., 2026): *E*_*r*_ = 0.79*eV, E*_*a*_ = 0.56*eV, E*_*d*_ = 0.49*eV*, and *E*_*e*_ = 0*eV*, which is why the conversion efficiencies are not temperature-dependent in the model description above. While the exact value of these activation energies is continuously updated, there is qualitative consensus that both consumption rates and death rates scale less quickly with temperature than population growth rates (Brown et al., 2004; Rall et al., 2012), while conversion efficiencies are largely temperature-independent (but see Lang et al., 2017).

The only activation energy we do not parametrize empirically but explore qualitatively in our analysis is *E*_*α*_, the temperature dependence of resource self-regulation. The temperature dependence of this parameter (and inversely, of the equilibrium density) remains poorly understood (Stockseth et al., 2025). We test three scenarios, illustrated in the top half of Fig. 1B:

1. *E*_*α*_ *< E*_*r*_: This scenario implies that the equilibrium density of the resource increases with warming, at least over the rising part of the thermal performance curves. There is theoretical and empirical evidence for this scenario (Lemoine, 2019; Stockseth et al., 2025; Vinton and Vasseur, 2022; Weiss et al., 2025). We use *E*_*α*_ = *E*_*a*_ = 0.56*eV* because consumer-resource theory predicts that self-regulation rates should scale proportionally to consumption rates (Fronhofer et al., 2024).
2. *E*_*α*_ = *E*_*r*_ = 0.79*eV* : This scenario implies that the equilibrium density of the resource stays constant with warming. There is some empirical evidence for this scenario, mostly from bacterial systems (Jarvis et al., 2016; Singh et al., 2011; Zwietering et al., 1991).
3. *E*_*α*_ *> E*_*r*_: This scenario implies that the equilibrium density of the resource decreases with warming, and has found empirical support mostly in phytoplankton systems (Bernhardt et al., 2018; Fussmann et al., 2014; West and Post, 2016). We use *E*_*α*_ = 1.44*eV* because of the canonical scaling of the equilibrium density/carrying capacity *K* as *E*_*K*_ = *−*0.65*eV* (Savage et al., 2004; Davis et al., 2026), and 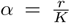, which implies *E*_*α*_ = *E*_*r*_ *− E*_*K*_ = 0.79*eV* + 0.65*eV* = 1.44*eV* .

### Analyses and parameters

We analyze the model first by deriving mutual invasibility criteria (Supporting Information S1), that is, we quantify whether a species can grow from low densities when the other species are present at equilibrium (Chesson, 2000; Amarasekare, 2008). We also use numerical simulations of the deterministic equations, carried out with the deSolve package in R (Soetaert et al., 2010), and stochastic simulations of the same equations for the early warning signal analysis later on. We implement demographic stochasticity with a bespoke modified Gillespie algorithm (Allen and Dytham, 2009). A Gillespie algorithm uses the master equation corresponding to the deterministic system to draw individual demographic events at rates emerging from the interplay of the underlying parameters (Black and McKane, 2012; Kokko, 2024). In all results shown below, we let simulations run for *t* = 25, 000 time steps. From the deterministic simulations, we infer equilibrium population sizes, stability of equilibria, and eigenvalues using the rootSolve package in R (Soetaert and Herman, 2009). From the stochastic simulations, we infer equilibrium population sizes by averaging across the last 1,000 time steps per replicate, and calculating medians and interquartile range across *n* = 100 replicates. We infer the dominant stochastic community state by counting the most frequent community outcome at the end of the simulations.

Across our analysis, we vary the temperature *T* as our most basic parameter (from 0^*°*^*C* to 40^*°*^*C*). To assess the relative importance of competitive vs. predatory intraguild processes, we mainly vary the activation energy of resource self-regulation (*E*_*α*_) as described above, as well as the relative strength of resource top-down regulation compared to resource self-regulation (see the lower half of Fig. 1B for an illustration). We keep death rates and conversion efficiencies constant and equal for consumer and predator. In sensitivity analyses, we vary the strength of intraguild competition, the strength of intraguild predation, baseline resource growth rates, and self-regulation rates. Supporting Information S2 summarizes all parameters and their tested values.

### Early warning signals

We use standard methods from early warning signal theory to predict warming-induced shifts in relative consumer and predator dominance in our system (see Box 1 for an explanation of the concept and its applicability in our system). In particular, we analyze the real parts of the leading eigenvalues of the deterministic system at equilibrium to assess whether the system would be subjected to critical slowing down (Kéfi et al., 2012; Patterson et al., 2021; Scheffer et al., 2009). We then calculate the coefficient of variation (CV), the skewness and the first-order autoregressive coefficient (AR1) (Dakos et al., 2012a) of each replicate stochastic time series, and compute medians and interquartile ranges of each indicator across replicate runs. In the results presented below, both skewness and AR1 are in absolute values. For these analyses, we subsample the stochastic time series at a frequency *s*. We test different sampling efforts by varying *s*, as different statistical indicators tend to have different requirements on the temporal resolution and length of time series (Dakos et al., 2012a; Weinans et al., 2019).

While all of the scenarios we test below converge to stable equilibria, some exhibit dampening oscillations depending on the initial conditions of the system. Such transients can continue to oscillate when demographic stochasticity is added to the system (Hastings et al., 2018). In order to avoid biasing the statistical indicators with artefacts generated by the initial conditions, we run separate stochastic simulations for our early warning signal analysis and start them from the deterministic equilibrium population sizes. In sensitivity analyses where this is not possible (because one of the species has an equilibrium population size of 0), we perform the early warning signal analysis after a burn-in of 10,000 time steps.

Our stochastic model can also capture environmental stochasticity in the form of a white noise with standard deviation *σ* around the mean temperature of the simulation drawn every *w* time units, which updates the rates of the system in real time. We test the effect of the presence, strength, and frequency of this thermal stochasticity in sensitivity analyses.

#### Box 1.

**Early warning signals and their application to our system**

Early warning signals occur as a result of “critical slowing down” as a system approaches a bifurcation. In mathematical terms, as the leading eigenvalue of the system approaches zero, the system is rendered more vulnerable to perturbations, and thus, indicators like variance, skewness, and autocorrelation of observable time series increase (Scheffer et al., 2009). This body of theory has traditionally been developed for catastrophic shifts at fold bifurcations, but has been shown to work for many other types of bifurcations (Kéfi et al., 2012). Fig. 1C illustrates how a time series of the consumer and its early warning signals should canonically respond to a change in temperature.

The appeal of early warning signals is that systemic changes, including catastrophic collapses, can be predicted based only on statistical attributes of a time series of a given observable. There are wide-ranging applications for these indicators, including climate tipping points, epileptic seizures, and stock market collapses (Scheffer et al., 2009). In ecology, they have been successfully applied from single population collapses (Dai et al., 2012) up to lake ecosystems close to eutrophication (Carpenter et al., 2011).

This potential simplicity of prediction stands in stark contrast with attempting to quantitatively predict collapses using mechanistic models. Forecasts based on the model described in Eqs. 1 and 2 would require estimation of 10 parameters and their thermal sensitivities – despite our choice of the simplest functional forms for both consumption rate and temperature dependence. Reliable estimation of all these parameters is perhaps possible in laboratory settings with fast-paced model organisms, but fails readily in the real world due to imprecision, confounding variables, and the massive sampling effort required. For these reasons, we test the reliability of several standard early warning signals to predict the warming-induced switches in dominance we expect in our system.

The key assumptions to validate in order to sensibly interpret early warning signals are (1) the stability of the incumbent equilibrium and (2) the presence of critical slowing down (Boettiger et al., 2013; Patterson et al., 2021). We verify whether our system fulfils these criteria by evaluating the eigenvalues of the Jacobian matrix of our system at equilibrium across the conditions explored in the analysis above. Indeed, all eigenvalues have negative real parts across the tested scenarios, indicating stable equilibria, and approach the zero at every switch described above (Supporting Information S4).

Early warning signals can fail for various reasons. External and correlated noise in the data, as well as low sampling resolution can render some indicators unreliable (Dakos et al., 2012b; Perretti and Munch, 2012; O’Regan and Burton, 2018). In more complex biotic contexts, perturbations do not necessarily align with the direction of the leading eigenvector of the system and therefore do not produce the phenomenon of critical slowing down (Boerlijst et al., 2013; Patterson et al., 2021). To test whether early warnings potentially work in our system, we run two-species control simulations, simulating only the interaction between either consumer or predator and the resource. Increases in CV consistently predict looming extinctions in both consumer and predator across scenarios and subsampling regimes (Supporting Information S5B-D). Increases in skewness also predict extinctions but require higher subsampling frequencies of the time series (Supporting Information S5E-G). Increases in AR1 are visible for some extinctions, but false positives occur even at high subsampling frequencies. Additionally, there is a strong interaction with the subsampling regime: In some settings, the autoregressive coefficient oscillates across temperatures due to an interaction with the stochastic oscillations of the system (Supporting Information S5H-J). From these control simulations, we conclude that CV represents the most reliable indicator for our system, followed by skewness, while AR1 is unreliable.

## Results

### Invasion analysis

The mutual invasibility criteria (see Supporting Information S1 for details) of consumer (*I*_*C*_) and predator (*I*_*P*_ ) at a given temperature are:

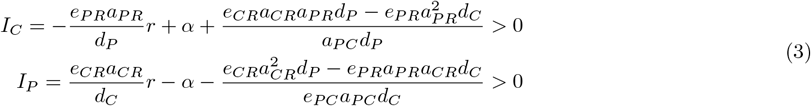

The invasibility criteria are expressed as composites of the pre-invasion dynamics of the resource and the incumbent species (first and second term), as well as an intraguild trade-off between conversion efficiencies, consumption rates, and death rates (third term). We have deliberately isolated the resource’s self-regulation rate *α* in the second term in each of these equations to understand better how the criteria as a whole change with temperature. This second term scales at a rate of *E*_*α*_, whereas the other terms have a composite thermal sensitivity: The first term consists of a constant (*e*_*ij*_), multiplied with a parameter that scales with *E*_*a*_ (*a*_*ij*_), multiplied with a parameter that scales with *E*_*r*_ (*r*), divided by a parameter that scales with *E*_*d*_ (*d*_*i*_). Taken together, the whole first term therefore scales with temperature at a rate of *E*_*a*_ + *E*_*r*_ *− E*_*d*_. By the same logic, the third term consists of two terms that both scale with temperature a rate of *E*_*a*_ + *E*_*a*_ + *E*_*d*_ *− E*_*a*_ *− E*_*d*_ = *E*_*a*_. We can therefore formulate two basic intuitions about the system and visualize them by plotting the dynamics of the invasibility criteria across temperatures (Fig. 2):

**Figure 2:**
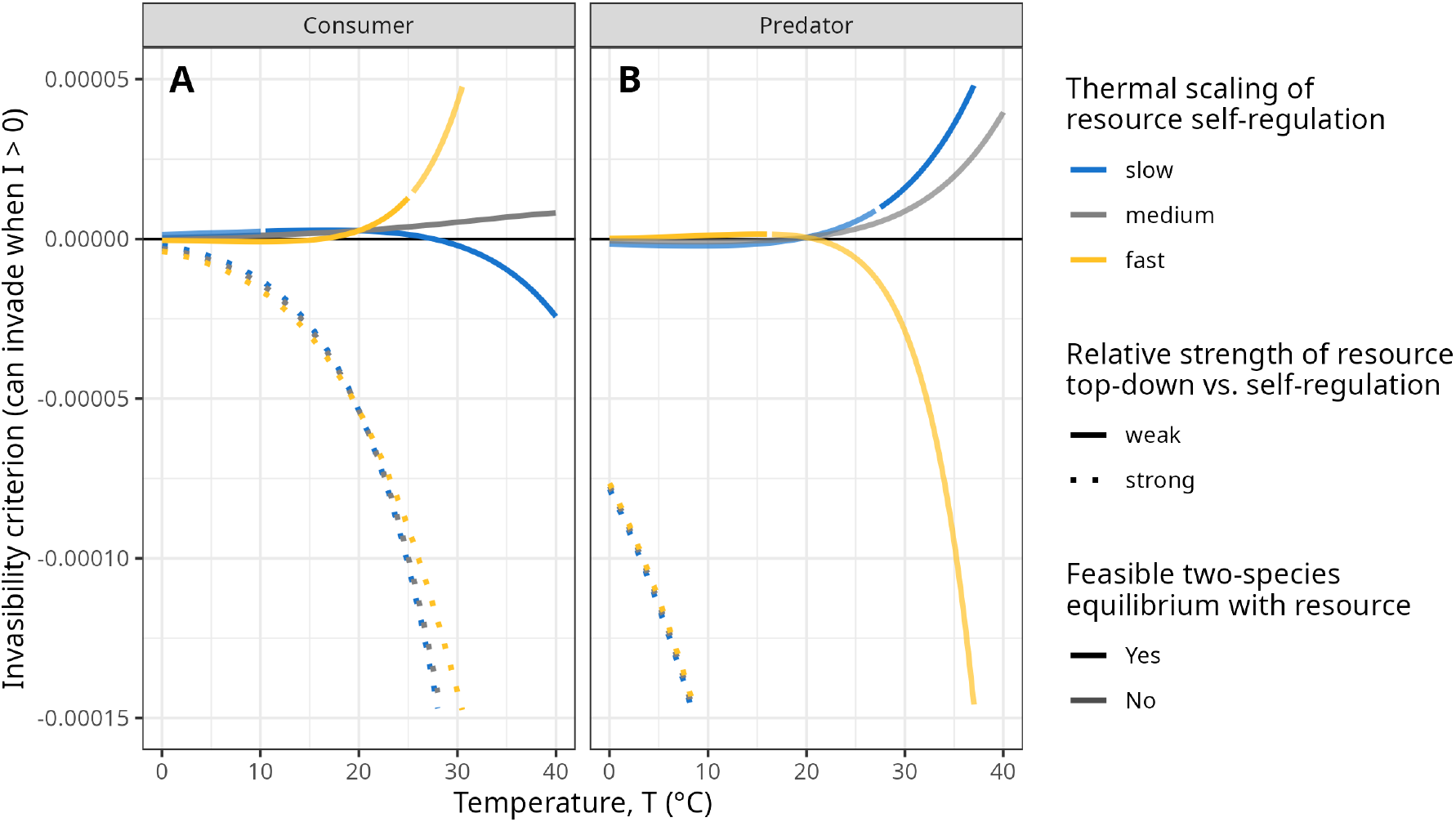
Invasibility criteria across temperatures for both consumer (A) and predator (B), differentiated by different activation energies of resource self-regulation *Eα* (colors; slow: *Eα* = 0.56, medium: *Eα* = 0.79, fast: *Eα* = 1.44) and resource top-down vs. self-regulation *β* (linetype; weak: *β* = 3, strong: *β* = 30). The feasibility of a nonzero two-species equilibrium between the focal species and the resource is displayed by the transparency. Other parameter settings are: *r*(*T*_0_) = 0.1, *α*(*T*_0_) = 0.00001, *γ* = 0.5, *δ* = 10, *e* = 0.05, *d*(*T*_0_) = 0.01.

1. The invasibility of either species can qualitatively change with temperature depending on the differential scaling of the different terms. As expressed above, the invasibility criteria show that the nature of the change depends on how quickly *α* scales compared to the other terms. When *α* scales relatively quickly with temperature, the consumer’s invasibility increases with warming, while the predator’s invasibility decreases (solid yellow lines in Fig. 2). The reverse is true when *α* scales relatively slowly with temperature (solid blue lines in Fig. 2). In both cases, the invasibility of both species flips from negative to positive or vice versa as temperatures increase.
2. The rate at which *α* scales with temperature only matters if *α* is of the same order of magnitude as the other parameters: In a strongly top-down regulated system (*a*_*ij*_ *>>> α*), the temperature dependence of the other terms dominates the evolution of the invasibility criteria. Indeed, when the resource is strongly top-down regulated (dotted lines in Fig. 2), there is no effect of the thermal scaling of resource self-regulation on invasibility. Both consumer and predator can persist in a two-species equilibrium with the resource across temperatures, but cannot invade each other, yielding a large range of bistability.

### Numerical simulations

Next, we conducted numerical simulations in which we started both consumer and predator from low densities. Given the large potential for bistability in the system shown above, these simulations do not aim to explore the outcomes of all possible initial conditions. Rather, they serve to illustrate the general phenomenon of temperature-induced switches in the dominance of consumer and predator.

The simulations (Fig. 3) confirm the intuitions provided by the analytical criteria. Qualitative switches in the dominance of consumer and predator occur as temperature changes. The direction of these switches depends on the thermal scaling of resource self-regulation, and on the relative strength of top-down regulation. We can discern the following cases:

**Figure 3:**
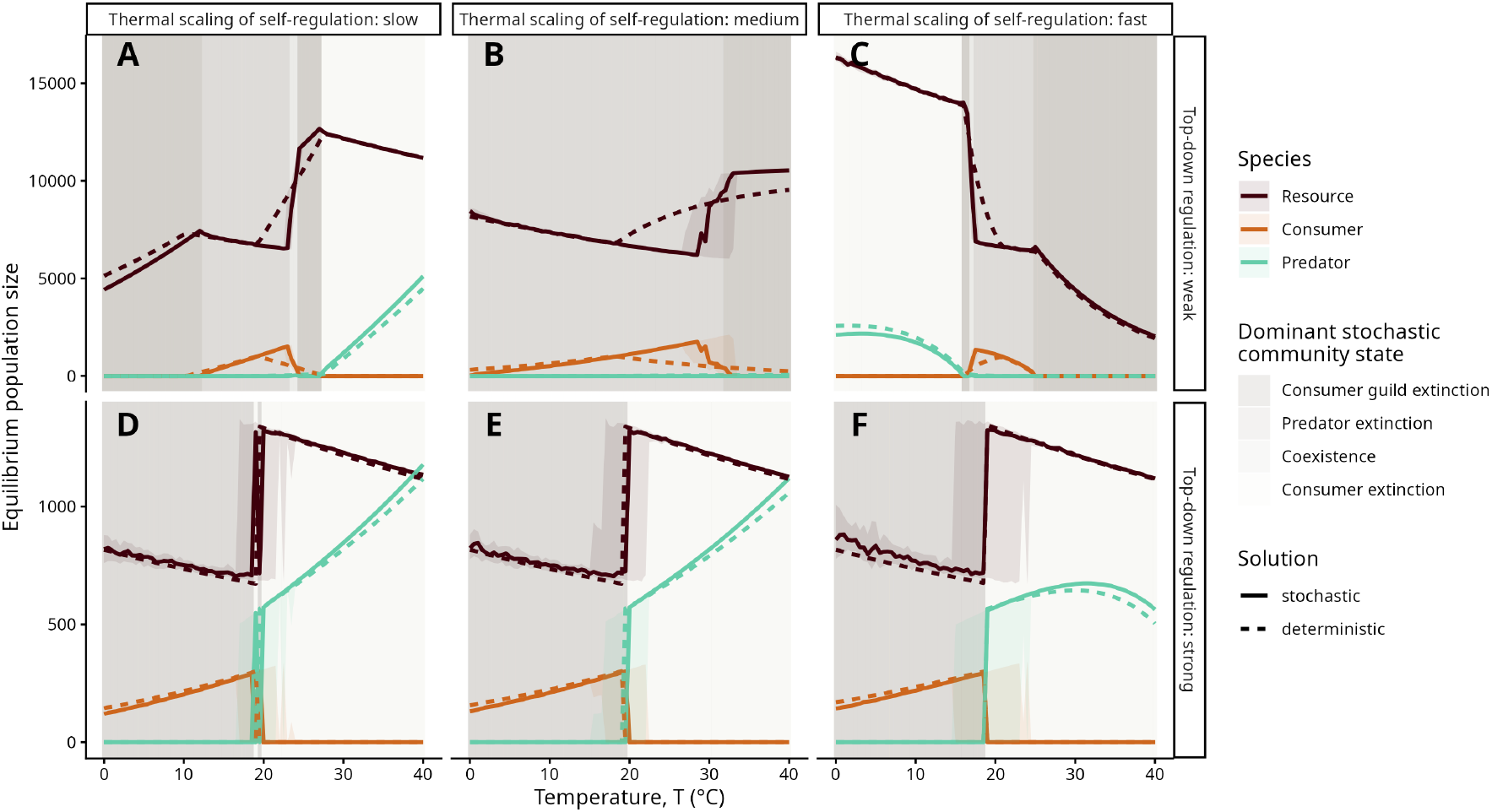
Equilibrium population densities of all species (colors) in the food web across temperatures (local x-axes), activation energies of resource self-regulation *Eα* (global x-axis; slow: *Eα* = 0.56, medium: *Eα* = 0.79, fast: *Eα* = 1.44), and resource top-down vs. self-regulation *β* (global y-axis; weak: *β* = 3, strong: *β* = 30). Numerical simulations of the deterministic system are in dashed lines; solid lines and shaded regions represent the median and interquartile range across 100 replicate runs of the stochastic simulations. Background is shaded according to the most frequent community outcome in the stochastic simulations. Other parameter settings are: *r*(*T*_0_) = 0.1, *α*(*T*_0_) = 0.00001, *γ* = 0.5, *δ* = 10, *e* = 0.05, *d*(*T*_0_) = 0.01. Initial population densities were *R* = 5000, *C* = 100, *P* = 50 in A-C, and *R* = 1000, *C* = 100, *P* = 50 in D-F to minimize overly large initial population oscillations. Equilibrium population densities were assessed at *t* = 25000. There was no additional thermal stochasticity.

1. When top-down regulation is relatively weak and resource self-regulation scales comparatively slowly (Fig. 3A), the system switches from extinction of both consumer and predator to predator extinction, then coexistence, then consumer extinction as temperature increases. The mechanism behind this is resource enrichment with temperature: Because of the slow scaling of self-regulation, the equilibrium density of the resource increases with temperature and shifts the dominant dynamic of the system from competitive to predatory.
2. When top-down regulation is relatively weak and resource self-regulation scales comparatively quickly (Fig. 3B,C), the succession of equilibrium states across temperature is now reversed. The resource contraction resulting from the faster scaling of *α* compared to *r* constrains the system as it warms, and moves it from a predatory to a competitive dynamic until neither consumer nor predator can be sustained.
3. When top-down regulation is relatively strong (Fig. 3D-F), the thermal scaling of resource self-regulation has no effect, and we observe a consistent switch from competitive to predatory dynamics with warming. In this case, the mechanism is different: Because the resource is so heavily top-down controlled (notice the difference in resource equilibrium densities between the upper and lower row of Fig. 3), it never reaches a density at which self-regulation becomes a meaningful factor. Instead, its maximum growth rate *r* and the increase of this rate with temperature drive enrichment of the system as it warms.

The results are robust to changes in baseline resource growth and self-regulation rates, as well as the strength of intraguild competition, intraguild predation, and resource top-down regulation (Supporting Information S3). We parametrized the model so that the switches occur around mild temperatures. Their exact location depends on the values of the underlying rates and the initial conditions. Introducing demographic stochasticity (compare dashed to solid lines in Fig. 3) does not change the outcome qualitatively, but renders the switches more diffuse. This is particularly evident in the scenarios with strong top-down control (Fig. 3D-F), where the potential of the system for bistability together with demographic stochasticity produce alternative stable states across a range of temperatures.

We note for the applied discussion below that a warming-induced switch from consumer to predator dominance always incurs a demographic release of the resource (brown lines in Fig. 3), as the predator is the weaker competitor for the resource.

### Early warning signals

We now turn to predicting the observed switches in consumer and predator dominance with generic early warning signals (see Box 1). In our analysis, we consider only replicate runs in which all three species stably coexist; an extinction of either consumer or predator reverts the analysis of early warning signals back to the two-species case (Box 1, Supporting Information 5-6). This filter restricts us to scenarios in which top-down regulation is weak, as there is no coexistence region between consumer and predator under strong top-down regulation. This is a trivial, but first result: Early warning signals preceding a switch in dominance between consumer and predator only work when the two species can stably coexist before the switch. They are therefore also not able to predict warming-induced switches in invasibility, as two-species equilibria between an incumbent and the resource do not exhibit critical slowing down at temperatures at which the invasibility of a potential invader becomes positive (compare Supporting Information S5 to the switchpoints in Fig. 2) without the introduction of the invader.

Fig. 4 shows the change in early warning signals across temperatures for the scenarios and replicate runs in which there is coexistence in the stochastic simulations. As in the two-species scenarios (Supporting Information S5), CV reliably predicts all extinctions, regardless of sampling frequency (Fig. 4A,B). Likewise, skewness is a good indicator of these extinctions, unless the sampling frequency gets too coarse (Fig. 4C,D, compare dashed lines to others). However, AR1 is not a reliable indicator of these extinctions, and different sampling frequencies can even revert the direction of the indicator trend (e.g., blue lines in Fig. 4E,F). The behavior of AR1 is remarkably similar across species (including the resource, see Supporting Information S7G-I). Upon inspection, the species’ population dynamics – despite starting from the equilibrium densities of a stable deterministic equilibrium – are forced into interdependent oscillations by stochastic deviations from the deterministic equilibrium (Supporting Information S8). This leads us to believe that the correlation between the different species’ population densities ‘overwrites’ the underlying autocorrelation structure and makes this indicator unreliable in our system. These results are robust to the presence, level, and frequency of thermal white noise (Supporting Information S9).

**Figure 4:**
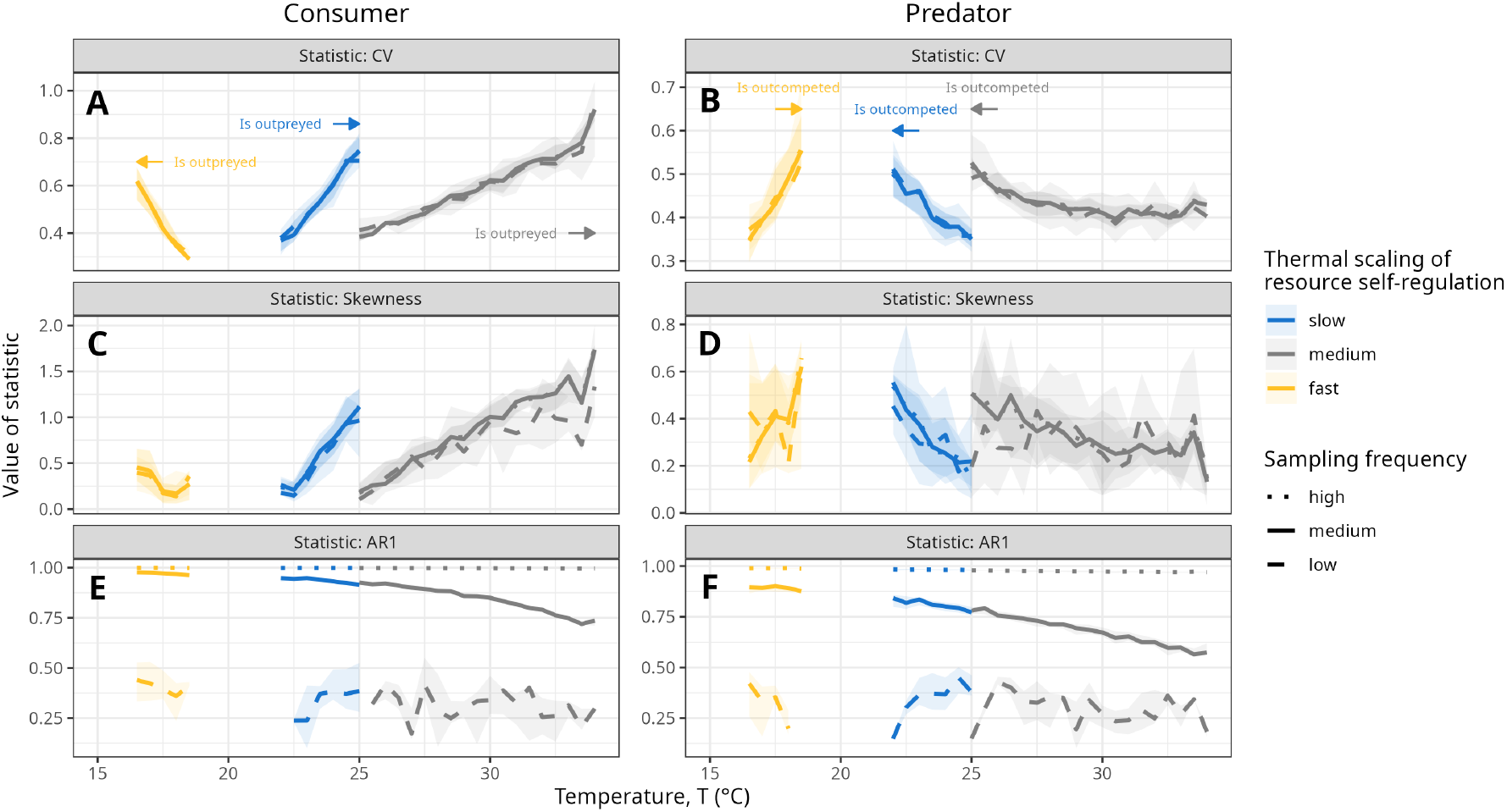
Early warning signals (rows) calculated from all stochastic time series in which there was three-species coexistence at *t* = 25000 for both consumer and predator (columns), temperatures (local x-axes), activation energies of resource self-regulation *E*_*α*_ (colors, values as above), and subsampling frequencies (linetype; high: *s* = 10, medium: *s* = 100, low: *s* = 1000 time steps). Lines and shaded regions represent the median and interquartile range across 100 replicate runs of the stochastic simulations. Arrows and annotations indicate the direction in which consumer or predator go extinct and why. Skewness and AR1 are expressed in absolute values. Initial population densities were set at the scenario-specific deterministic equilibrium density. There was no thermal stochasticity. Other parameter settings are: *r*(*T*_0_) = 0.1, *α*(*T*_0_) = 0.00001, *β* = 3, *γ* = 0.5, *δ* = 10, *e* = 0.05, *d*(*T*_0_) = 0.01.

## Discussion

By exploring the temperature dependence of a simple IGP system, we have shown that empirically established differences between the thermal sensitivity of rates on the resource level and on higher trophic levels can produce qualitative shifts in invasion and equilibrium outcomes across temperatures. These shifts are driven by temperature-induced changes in resource enrichment, which in turn changes the relative importance of competitive vs. predatory intraguild processes.

The few previous empirical studies on the effects of temperature on IGP systems have all found increased consumption rates with warming (Sentis et al., 2014; Frances and McCauley, 2018; Rogers et al., 2018). They predicted either system destabilization or predator dominance at higher temperatures. However, in these systems (ladybugs, dragonflies, blue/green crabs), it is not feasible to assess long-term dynamics of the resource without large efforts, which might lead to biased predictions (Briggs and Borer, 2005). Our model shows that, despite an increase in intraguild predation rates, both consumer exclusion and predator exclusion are possible outcomes as temperatures increase. The outcome depends on the two key parameters we varied in our model: the temperature dependence of resource self-regulation, and the relative strength of resource self-vs. top-down regulation. We have kept our model as minimal as possible, and the mechanism of an enrichment-mediated shift in intraguild power dynamics holds also for more complex functional response types (Mylius et al., 2001). Of course, other factors and complexities such as ontogenetic diet shifts (Thunell et al., 2021) or thermal asymmetries (Gibert et al., 2022) can change these outcomes qualitatively.

Our model results therefore prompt two rather fundamental questions: (1) Is there a general temperature dependence of resource self-regulation, and (2) are IGP systems rather bottom-up or top-down controlled?

The temperature dependence of resource self-regulation remains an open empirical question (Stockseth et al., 2025). Only a few studies have explicitly assessed intraspecific competition as a focal parameter (as opposed to equilibrium density as the collective property). These studies have mostly found some sort of unimodality in the response to temperature (DeLong and Lyon, 2020; Johnson et al., 2015; Stockseth et al., 2025). Yet, thermal performance curves or scaling relationships over the rising part of these curves have not been estimated. In terms of equilibrium density, some experiments with phytoplankton (Bernhardt et al., 2018; West and Post, 2016) and meta-analyses seem to suggest a rather strong negative temperature sensitivity (Fussmann et al., 2014; Davis et al., 2026), while another meta-analysis on phytoplankton and insects reported a majority of unimodal responses to temperature (Lemoine, 2019). Other results show constant equilibrium densities across temperatures (Jarvis et al., 2016; Singh et al., 2011; Zwietering et al., 1991). The empirical evidence overall only partially agrees with theoretical predictions (Uszko et al., 2017; Lemoine, 2019; Vinton and Vasseur, 2022). Our results demonstrate how important a general and mechanistic understanding of the temperature-dependence of self-regulation is to predict the dynamics of food webs subject to warming.

Regarding bottom-up or top-down control of resources in IGP systems, we can resort to global meta-analyses. Results vary across taxa: While plants seem to be rather equally bottom-up and top-down controlled (Gruner et al., 2008), small mammals exhibit strong bottom-up control (Prevedello et al., 2013), whereas terrestrial insect herbivores seem to be rather top-down controlled (Vidal and Murphy, 2017). The extent of top-down vs. bottom-control varies within the same food web: Marine phytoplankton are rather bottom-up controlled, whereas piscivorous fish in the same community are rather top-down controlled (Lynam et al., 2017). Warming seems to have mixed effects on these relationships (Rodríguez-Castañeda, 2012; Vidal et al., 2026). Given the variety of patterns found across taxa, we would reason that all scenarios explored above are equally biologically plausible. For example, we would expect that in IGP modules in which marine phytoplankton is the basal resource (i.e., weak top-down control and strong thermal sensitivity of self-regulation), warming would increase consumer dominance over the predator. Conversely, in IGP modules in which herbivorous insects are the basal resource (i.e., strong top-down control), we would expect warming to increase predator dominance over the consumer.

We are not the first to stress the importance of the balance between top-down and bottom-up processes for the response of interacting species to warming. Several studies before us have shown that integrating the temperature dependence of a resource is key to predicting a consumer’s response to changes in temperature (e.g., Amarasekare, 2015; West and Post, 2016; Uszko et al., 2017; Vinton and Vasseur, 2022). Such integration can unveil surprising outcomes: Binzer et al. (2012) showed that increasing temperature can counteract the destabilizing effect of nutrient enrichment on food chains. Here, we have shown how the interaction between top-down and bottom-up processes can change the power structure of a food web as the system warms. Importantly, these changes occur not because of external, interspecific differences, e.g., in thermal optima or behavior, but only because of mechanisms contained within the most essential system-defining rates.

Our analysis focuses on a local IGP system. However, our results show that the same three species, without any differences in their thermal niches, could produce different community states across a landscape structured by a thermal gradient. Spatial dynamics can change local IGP equilibria (Bampfylde and Lewis, 2007), and this depends on possible intraguild asymmetries in dispersal (Amarasekare, 2006) and on the connectivity on the landscape (Okuyama, 2008).

Extending our model to include spatial structure would be an important next step to understand the effect of temperature on IGP systems more cohesively. A key ingredient for this extension would be a better understanding of the temperature dependence of dispersal, which is currently sorely lacking (Amarasekare, 2024).

### Prediction and application

We have tested the capacity of several generic statistical indicators to serve as early warning signals for the warming-induced switches in dominance. We found varying results for different indicators: Variance- and skewness-based indicators performed generally well, while first-order autocorrelation was not reliable. The sampling effort had a large influence on the results of skewness and autocorrelation. Our results align with those of Perretti and Munch (2012), who show that variance-based indicators are the most reliable of the standard indicators, especially under ecologically realistic levels of noise. We have not tested any autocorrelation structure in the noise, but their work suggests that these indicators can be robust even to pink or red noise. While it is promising that CV and skewness of a time series can predict the shifts in our system, skewness is known to produce false negatives from time to time (Dakos et al., 2012a) and suffers from high noise levels (Perretti and Munch, 2012), and variance-based indicators have been shown to be rather sensitive to certain types of environmental noise (Dakos et al., 2012b; O’Regan and Burton, 2018).

We trace the failure of AR1 to predict extinctions in our system to interdependent oscillations caused by stochastic deviations from a stable deterministic equilibrium. The capacity of stochastic systems to propagate transient deterministic oscillations is well-known (Hastings et al., 2018) and it is unsurprising that an intrinsic autocorrelation structure is reducing the capacity of an autocorrelation-based measure to predict transitions. However, our results add to the growing tally of studies that demonstrate the difficulty of successfully using early warning signals in more complex biotic contexts. Some ecosystem collapses do not show signs of critical slowing down if perturbations are not occurring in the direction of the leading eigenvector (Boerlijst et al., 2013). Consequently, there exist theoretical and practical guidelines for identifying early warning signals in multispecies communities (Dakos, 2018; Patterson et al., 2021; Weinans et al., 2019). However, the failure of early warning signals in these contexts is frequent and sometimes unavoidable. Therefore, we argue that a mechanistic understanding of the system in question remains crucial to predict qualitative shifts in invasibility and equilibrium outcomes under global change.

Our results have direct implications for agricultural applications. Many biological control methods in agricultural systems use insect natural enemies that can engage in some form of IGP (Polis and Holt, 1992; Briggs and Borer, 2005; Martin et al., 2013). Our results show that, depending on the system, biological control agents can become more or less effective with warming. In systems, in which top-down control is rather strong, the weaker competitor for the pest is more likely to persist at higher temperatures, increasing herbivore density and thus reducing pest control. This is likely the case for agricultural systems, in which aphids are often the herbivore pest, and top-down control is prevalent (Vidal and Murphy, 2017). In systems, in which bottom-up control is rather strong, a good understanding of the pest’s dynamics across temperatures without natural enemies is crucial. If the pest’s self-regulation increases rapidly with temperature, the stronger competitor for the pest is more likely to persist, and biological control will become more effective with warming. In the absence of such data, our analysis suggests that variance-based indicators are likely the most reliable to predict warming-induced shifts in real IGP systems. Of course, these statistical indicators only work for systems in which the species in question are both stably present; as reasoned above, generic early warning signals cannot predict shifts in invasibility. For these more devious shifts, a more mechanistic understanding of the system and the natural history of potential invaders is necessary.

### Conclusion

By focusing on a classic and tractable three-species food web module, we have revealed non-trivial, built-in shifts in the power dynamics of this module as the temperature changes. We have also shown that these shifts are not reliably predicted by all classical early warning signals. To reliably forecast small and large food webs in the face of anthropogenic climate change, we need a more general understanding of the temperature dependence of resource self-regulation, and scalable ways to quantify it in the field.

## Data Availability

Code to reproduce all results will be made publicly available via GitHub/Zenodo upon acceptance.

## Author contributions

P.K. and E.A.F. designed the research. P.K. wrote the code, performed and analyzed the simulations, and wrote the first draft. E.A.F. acquired funding, supervised the work, and reviewed and edited the draft.

## Acknowledgements

We thank Vasilis Dakos, Nathan Humbert, Sonia Kéfi, and Emilio Mora for helpful discussions. This work was supported by a grant from the Agence Nationale de la Recherche (ANR-23-CE02-0030). The authors declare no competing interests.

