## Supporting Information for "Warming-induced switches in dominance are built into intraguild predation systems"

June 18, 2026

#### Supporting Information S1: Derivation of mutual invasibility criteria

The model without temperature dependence writes:

$$\begin{aligned}\frac{1}{R}\frac{dR}{dt} &= r - \alpha R - a_{CR}C - a_{PR}P \\ \frac{1}{C}\frac{dC}{dt} &= e_{CR}a_{CR}R - a_{PC}P - d_C \\ \frac{1}{P}\frac{dP}{dt} &= e_{PR}a_{PR}R + e_{PC}a_{PC}C - d_P\end{aligned} \tag{1}$$

To derive mutual invasibility criteria for consumer and predator, we assume the resource is at nonzero equilibrium $R^* > 0$, at which

$$\frac{1}{R}\frac{dR}{dt} = r - \alpha R - a_{CR}C - a_{PR}P = 0 \tag{2}$$

which yields

$$R^* = \frac{r - a_{CR}C - a_{PR}P}{\alpha} \tag{3}$$

##### Invasibility criterion for predator

To derive an invasibility criterion for the predator, we start by deriving the expressions for the equilibrium densities of the system when $P = 0$. Equation (3) simplifies to:

$$R^* = \frac{r - a_{CR}C}{\alpha} \tag{4}$$

and, assuming $C^* > 0$, we have

$$\frac{1}{C}\frac{dC}{dt} = e_{CR}a_{CR}R - d_C = 0 \tag{5}$$

Inserting equation (4) into equation (5) yields:

$$e_{CR}a_{CR}\frac{r - a_{CR}C}{\alpha} - d_C = 0 \tag{6}$$

which simplifies to

$$C^* = \frac{r}{a_{CR}} - \frac{d_C\alpha}{e_{CR}a_{CR}^2} \tag{7}$$

Reinserting this expression into equation (4) yields:

$$R^* = \frac{r - a_{CR}(\frac{r}{a_{CR}} - \frac{d_C\alpha}{e_{CR}a_{CR}^2})}{\alpha} \tag{8}$$

which simplifies to

$$R^* = \frac{d_C}{e_{CR}a_{CR}} \tag{9}$$

For the predator to invade from low densities, it must have a positive growth rate when resource and consumer are at their equilibrium densities:

$$\frac{1}{P}\frac{dP}{dt} = e_{PR}a_{PR}R^* + e_{PC}a_{PC}C^* - d_P > 0 \tag{10}$$

Inserting our expressions from equations (7) and (9) yields:

$$e_{PR}a_{PR}\frac{d_C}{e_{CR}a_{CR}} + e_{PC}a_{PC}(\frac{r}{a_{CR}} - \frac{d_C\alpha}{e_{CR}a_{CR}^2}) - d_P > 0 \tag{11}$$

which simplifies to

$$e_{PR}a_{PR}a_{CR}d_C + e_{PC}a_{PC}e_{CR}a_{CR}r - e_{PC}a_{PC}d_C\alpha - e_{CR}a_{CR}^2d_P > 0 \tag{12}$$

We can isolate $\alpha$ and define the invasibility criterion:

$$I_P = \frac{e_{CR}a_{CR}}{d_C}r - \alpha - \frac{e_{CR}a_{CR}^2d_P - e_{PR}a_{PR}a_{CR}d_C}{e_{PC}a_{PC}d_C} > 0 \tag{13}$$

We have rearranged the third term to be positive in our setup where conversion efficiencies and death rates are equal across consumer and predator and the consumer is the stronger competitor for the resource by way of $a_{CR} > a_{PR}$.

**Invasibility criterion for consumer**

We follow the same procedure to derive an invasibility criterion for the consumer. The two-species equilibria for predator and resource are analogous to before:

$$P^* = \frac{r}{a_{PR}} - \frac{d_P\alpha}{e_{PR}a_{PR}^2} \tag{14}$$

and

$$R^* = \frac{d_P}{e_{PR}a_{PR}} \tag{15}$$

For the consumer to invade from low densities, the following inequality must hold:

$$\frac{1}{C}\frac{dC}{dt} = e_{CR}a_{CR}R^* - a_{PC}P^* - d_C > 0 \tag{16}$$

Inserting equations (14) and (15) into this inequality yields:

$$e_{CR}a_{CR}\frac{d_P}{e_{PR}a_{PR}} - a_{PC}(\frac{r}{a_{PR}} - \frac{d_P\alpha}{e_{PR}a_{PR}^2}) - d_C > 0 \tag{17}$$

which simplifies to

$$e_{CR}a_{CR}a_{PR}d_P - e_{PR}a_{PR}a_{PC}r + a_{PC}d_P\alpha - e_{PR}a_{PR}^2d_C > 0 \tag{18}$$

As above, we can isolate $\alpha$, rearrange for the third term to be positive, and define the invasibility criterion:

$$I_C = -\frac{e_{PR}a_{PR}}{d_P}r + \alpha + \frac{e_{CR}a_{CR}a_{PR}d_P - e_{PR}a_{PR}^2d_C}{a_{PC}d_P} > 0 \tag{19}$$

### Supporting Information S2: Parameter explanations and ranges

Table S2: Explored parameters and their ranges of values

| Parameter | Description | Explored (standard) values |
|---|---|---|
| $r$ | Resource maximum growth rate | 0.01, 0.04, 0.07, 0.1, 0.13, 0.16, 0.19 |
| $\alpha$ | Resource self-regulation rate | 0.000001, 0.00001 |
| $e_{ij}$ | Conversion efficiency of species $i$ when feeding on species $j$ | 0.05 |
| $a_{ij}$ | Consumption rate of species $i$ when feeding on species $j$ | Emerges from parameters $\alpha, \beta, \gamma, \delta$, see below |
| $d_i$ | Death rate of species $i$ | 0.01 |
| $T$ | Temperature (°C) | 0-40 in increments of 0.5 |
| $T_0$ | Baseline temperature (°C) | 20 |
| $k$ | Boltzmann constant ($\frac{eV}{K}$) | $8.617 \times 10^{-5}$ |
| $E_r$ | Activation energy of resource maximum growth rate (eV) | 0.79 |
| $E_a$ | Activation energy of consumer and predator consumption rate (eV) | 0.56 |
| $E_a$ | Activation energy of consumer and predator death rate (eV) | 0.49 |
| $E_\alpha$ | Activation energy of resource self-regulation (eV) | 0.56, 0.79, 1.44 |
| $\beta$ | Relative strength of resource top-down vs. self-regulation, such that $a_{CR}(T_0) = \beta\alpha(T_0)$ | 3,10,30 |
| $\gamma$ | Relative strength of intraguild predation, such that $a_{PR}(T_0) = \gamma\beta\alpha(T_0)$ | 3,10,30 |
| $\delta$ | Relative strength of intraguild competition, such that $a_{PC} = \delta\alpha(T_0)$ | 0.25,0.5,0.75 |

### Supporting Information S3: Sensitivity analysis for numerical simulations

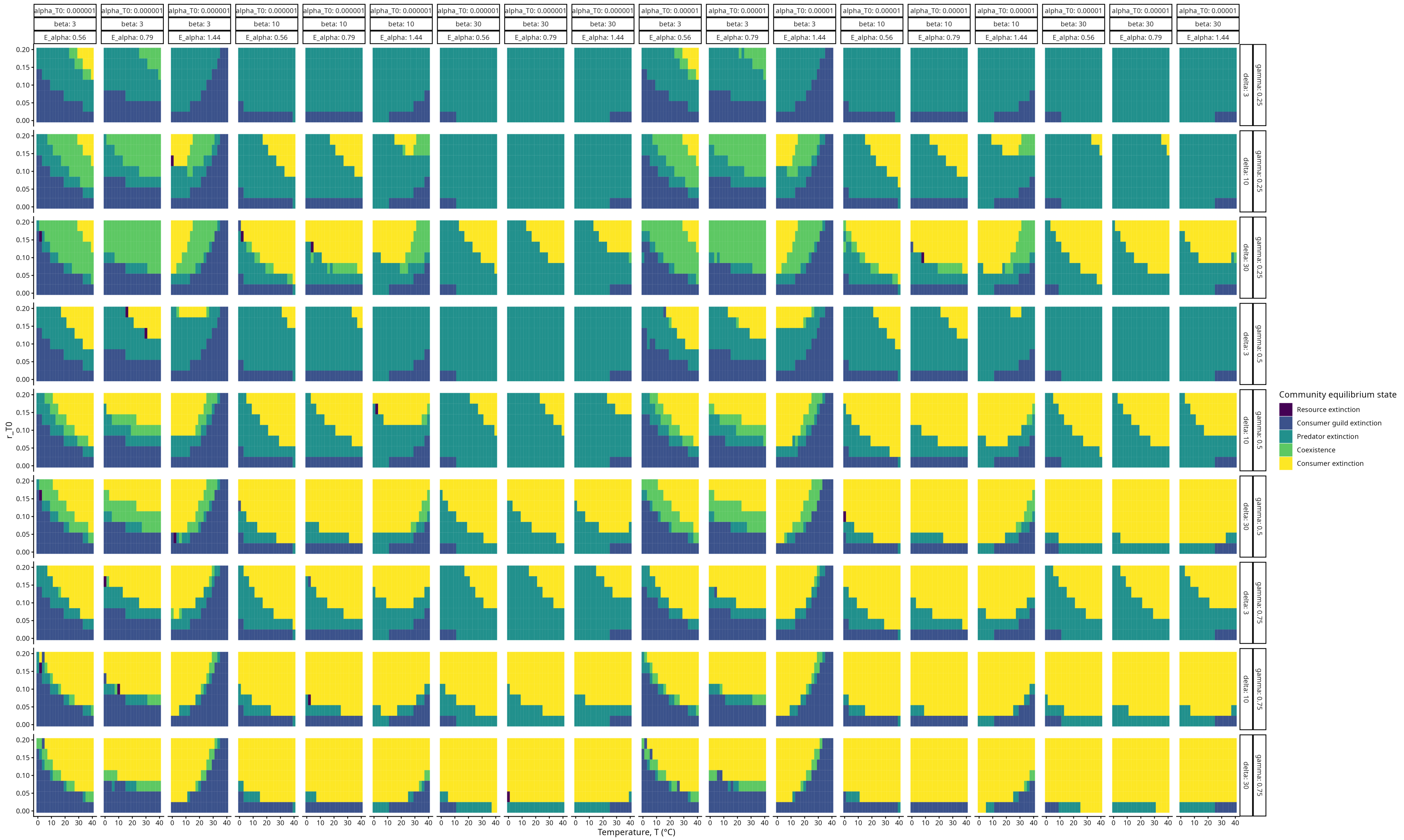


Figure S3: Sensitivity analysis for the numerical simulations. Colors in the heatmap encode the equilibrium community state at $t = 25000$ in the deterministic simulations. Species were considered extinct when their density was below $1e^{-5}$. Parameters varied are: Temperature (local x-axes), growth rate at baseline temperature $r(T_0)$ (local y-axes), resource self-regulation at baseline temperature $\alpha(T_0)$ (global x-axis, first level), relative strength of resource top-down vs. self-regulation $\beta$ (global x-axis, second level), activation energy of resource self-regulation $E_\alpha$ (global x-axis, third level), relative strength of intraguild competition $\gamma$ (global y-axis, first level), and relative strength of intraguild predation $\delta$ (global y-axis, second level). Initial population densities were $R = 2500, C = 100, P = 50$. Other parameter setting were $e = 0.05, d(T_0) = 0.01$.

### Supporting Information S4:
### Verifying assumptions for early warning signal analysis

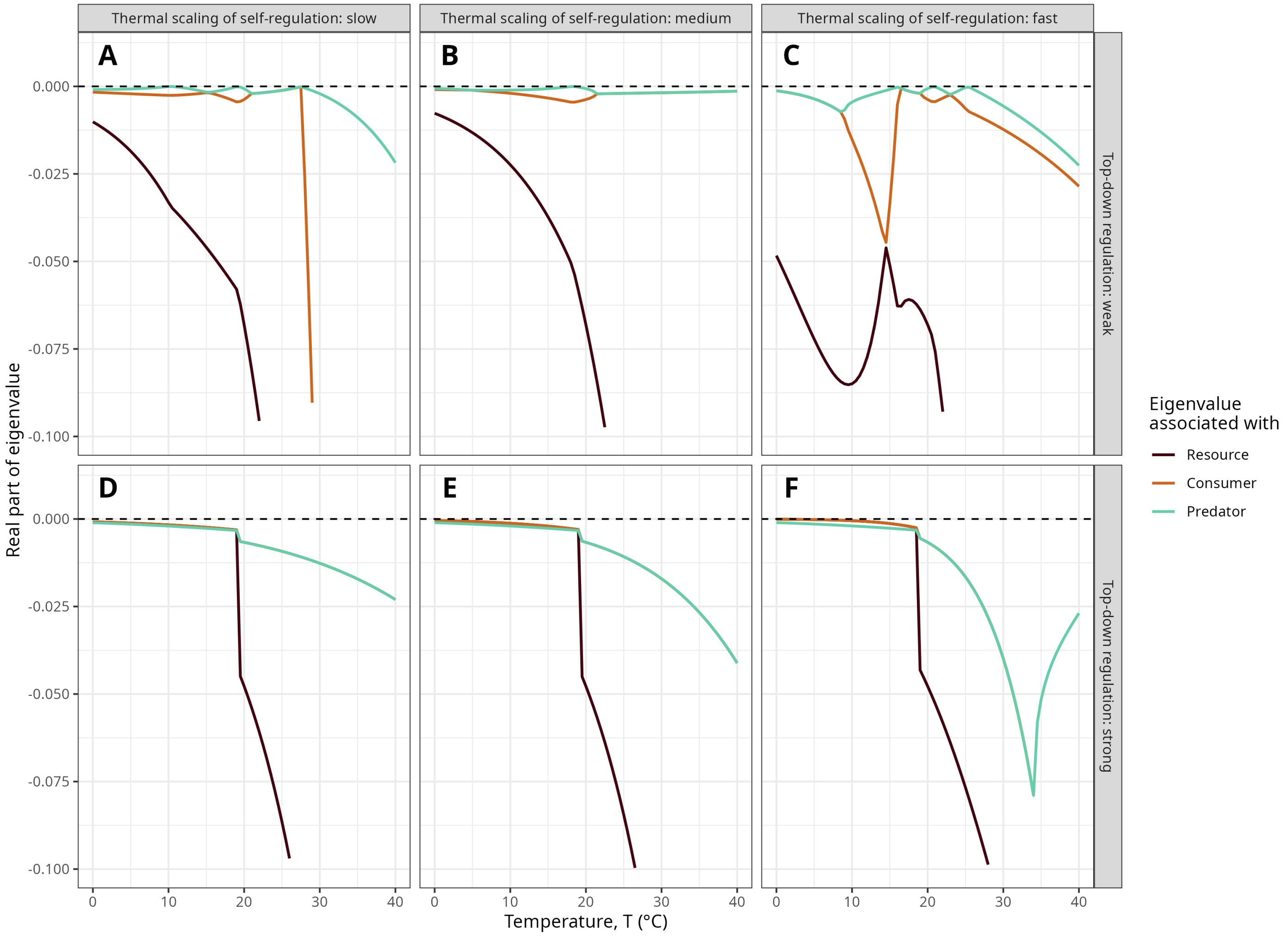


Figure S4: Real parts of the eigenvalues of the Jacobian at equilibrium associated with all species (colors) in the food web across temperatures (local x-axes), activation energies of resource self-regulation $E_\alpha$ (global x-axis; slow: $E_\alpha = 0.56$, medium: $E_\alpha = 0.79$, fast: $E_\alpha = 1.44$), and resource top-down vs. self-regulation $\beta$ (global y-axis; weak: $\beta = 3$, strong: $\beta = 30$). Other parameter settings are: $r(T_0) = 0.1, \alpha(T_0) = 0.00001, \gamma = 0.5, \delta = 10, e = 0.05, d(T_0) = 0.01$.

### Supporting Information S5: Control case for early warning signals

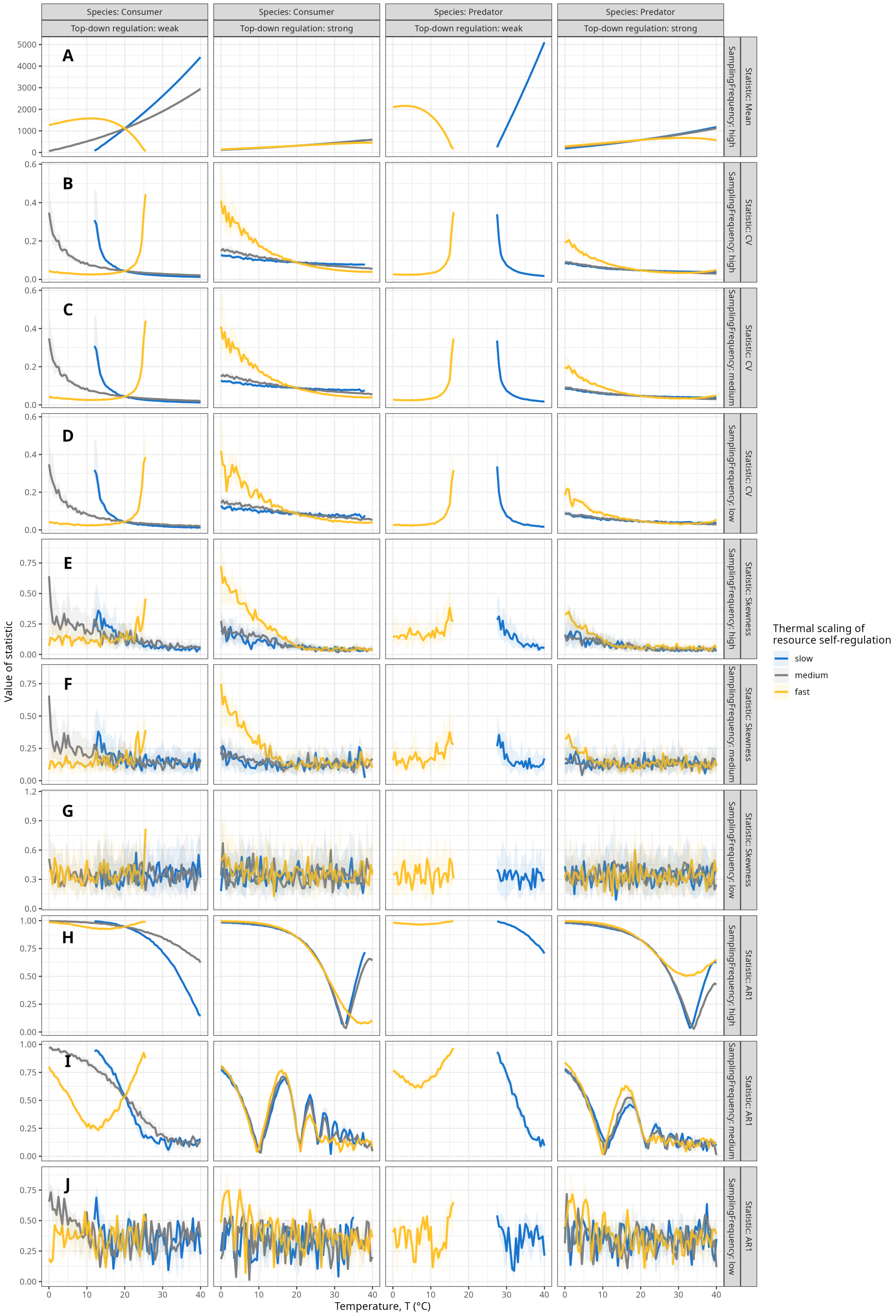


Figure S5: Early warning signals calculated from stochastic simulations in which only one of consumer or predator was started at nonzero densities with the resource. Early warning signals (global y-axis, first level) are shown for consumer and predator (global x-axis, first level), temperatures (local x-axes), activation energies of resource self-regulation $E_\alpha$ (colors, values as before), resource top-down vs. self-regulation $\beta$ (global x-axis, second level; values as before), and subsampling frequencies (global y-axis, second level; high: $s = 10$, medium: $s = 100$, low: $s = 1000$ time steps). The population mean is shown in the top row and in only one subsampling frequency to show where species go extinct. Early warning signals were calculated after a burn-in of 10,000 time steps. Lines and shaded regions represent the medians and interquartile ranges across 25 replicate runs of the stochastic simulations. Skewness and AR1 are expressed in absolute values. Initial population densities and other parameters are as in Figure 3.

### Supporting Information S6:
### Early warning signals for non-coexistence scenarios

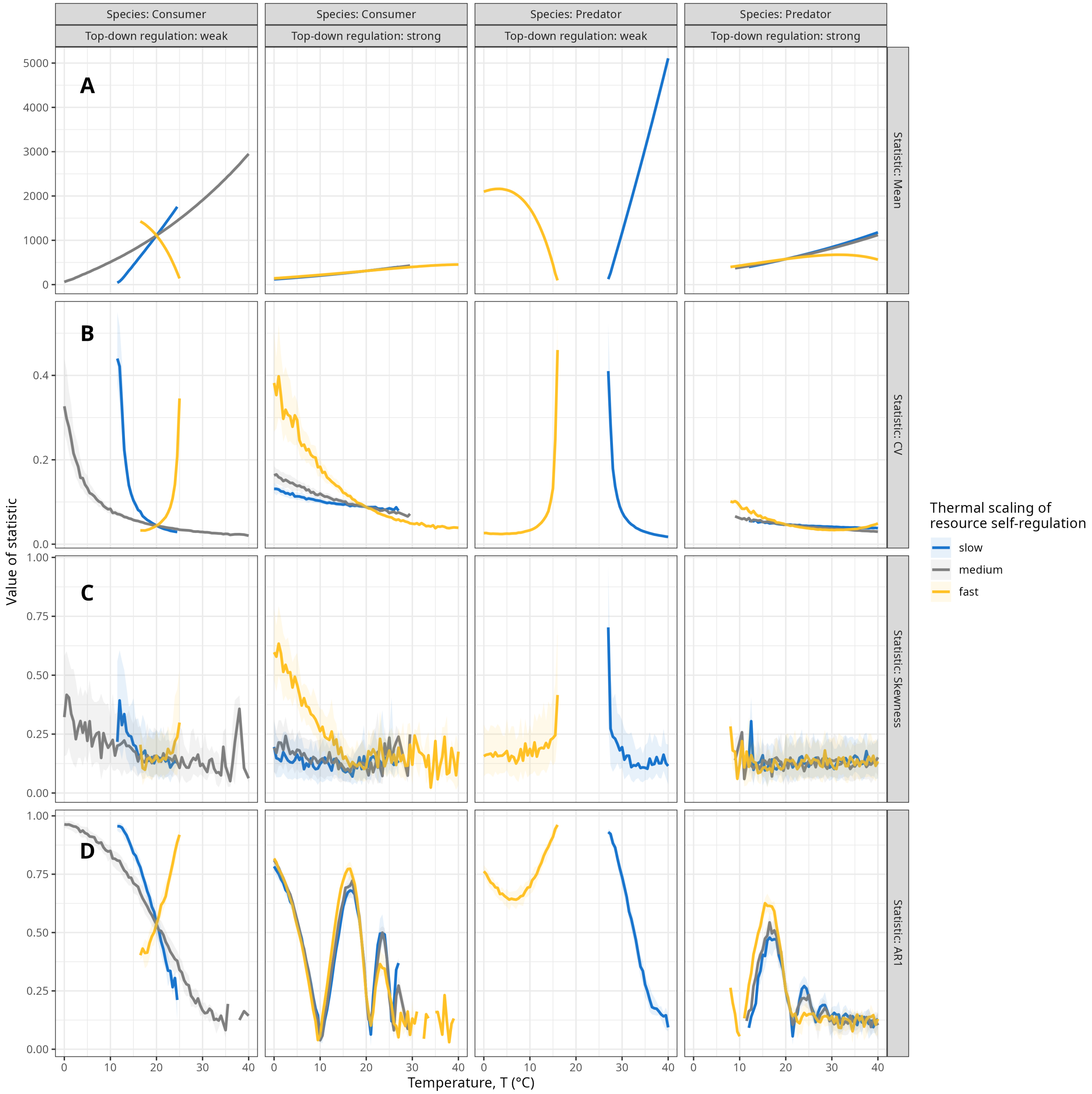


Figure S6: Early warning signals calculated from the stochastic simulations that produced Figure 3 but filtered for scenarios and replicate runs in which only one of consumer or predator survived throughout the entire length of the time series considered for the early warning signals. Early warning signals (global y-axis) are shown for consumer and predator (global x-axis, first level), temperatures (local x-axes), activation energies of resource self-regulation $E_\alpha$ (colors, values as before), and resource top-down vs. self-regulation $\beta$ (global x-axis, second level; values as before). The population mean is shown in the top row to demonstrate where species go extinct. Early warning signals were calculated after a burn-in of 10,000 time steps and are shown at a subsampling frequency of $s = 100$ time steps. Lines and shaded regions represent the medians and interquartile ranges across 100 replicate runs of the stochastic simulations. Skewness and AR1 are expressed in absolute values. Initial population densities and other parameters are as in Figure 3.

### Supporting Information S7: Early warning signals in the resource

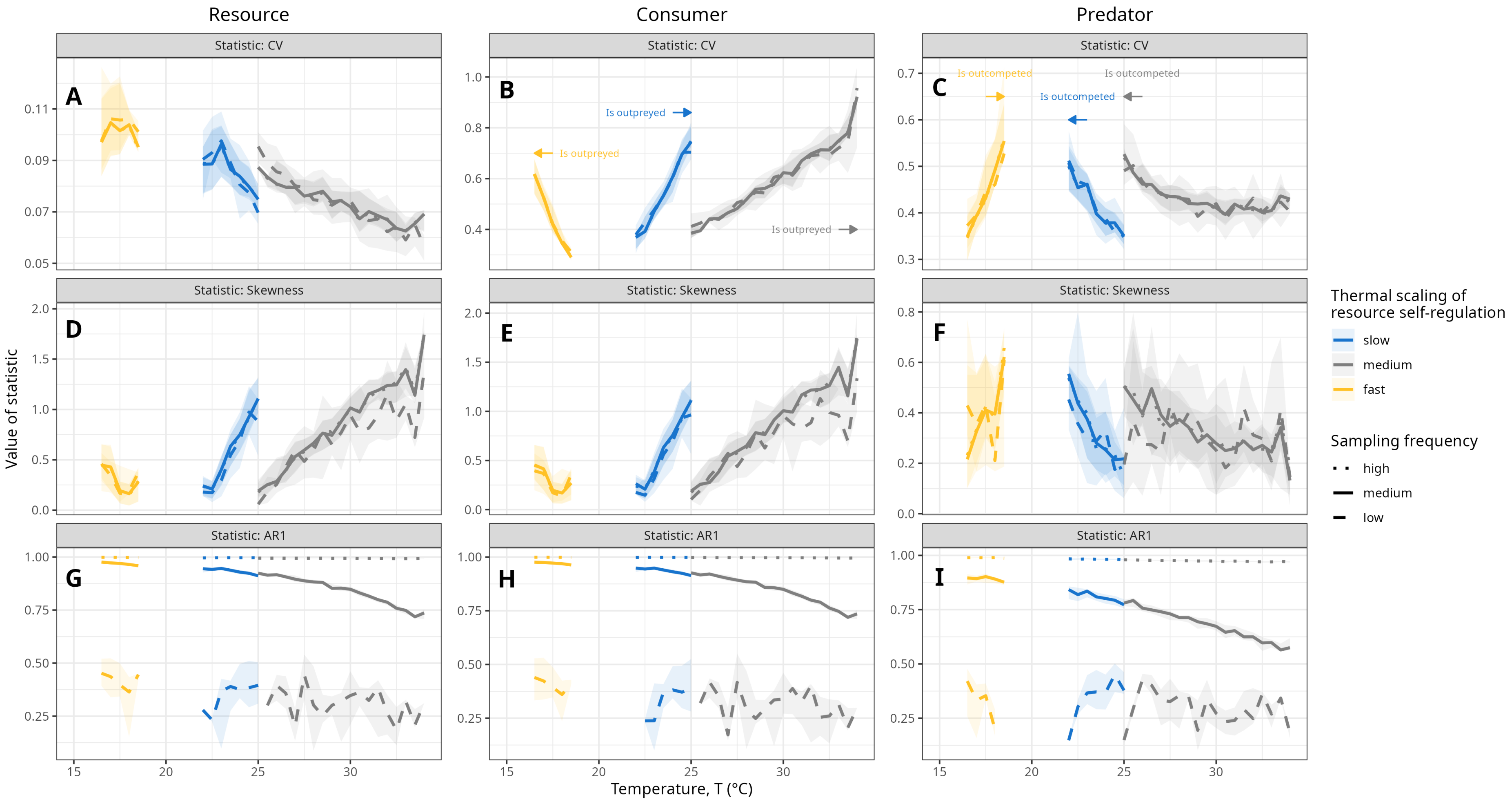


Figure S7: Early warning signals calculated from all stochastic time series in which there was three-species coexistence at $t = 25000$ across species (columns), temperatures (local x-axes), activation energies of resource self-regulation $E_\alpha$ (colors, values as above), and subsampling frequencies (linetype; high: $s = 10$, medium: $s = 100$, low: $s = 1000$ time steps). Lines and shaded regions represent the median and interquartile range across 100 replicate runs of the stochastic simulations. Arrows and annotations indicate the direction in which the respective species goes extinct and why. Skewness and AR1 are expressed in absolute values. Initial population densities were set at the scenario-specific deterministic equilibrium density. There was no thermal stochasticity. Other parameter settings are: $r(T_0) = 0.1, \alpha(T_0) = 0.00001, \beta = 3, \gamma = 0.5, \delta = 10, e = 0.05, d(T_0) = 0.01$.

### Supporting Information S8:
### Population cycling in the stochastic simulations

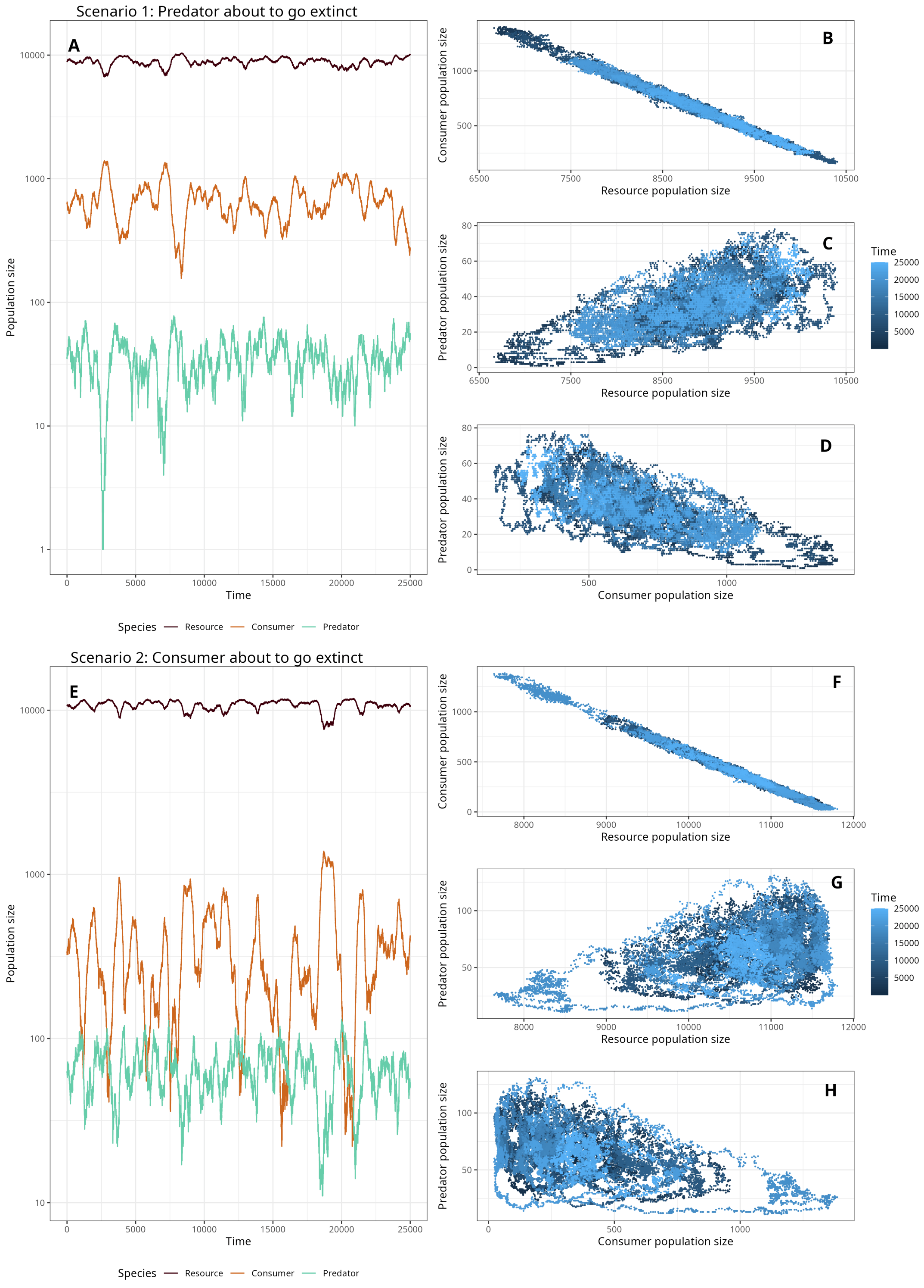


Figure S8: Representative replicate time series of all three species' population densities (left column) and pairwise correlations between species' population densities (right column). The top half represents a scenario in which the predator is about to go extinct if the environment becomes any colder, the bottom half a scenario in which the consumer is about to go extinct if the environment becomes any warmer. The scenario is taken from the extremes of the coexistence space at a slow scaling of resource self-regulation $E_\alpha = 0.56$, with the upper scenario at 22.5°C (replicate 13) and the lower scenario at 25°C (replicate 0). Initial population densities were set at the scenario-specific deterministic equilibrium density. Other parameter settings are: $r(T_0) = 0.1, \alpha(T_0) = 0.00001, \beta = 3, \gamma = 0.5, \delta = 10, e = 0.05, d(T_0) = 0.01$.

### Supporting Information S9: Early warning signals with thermal noise

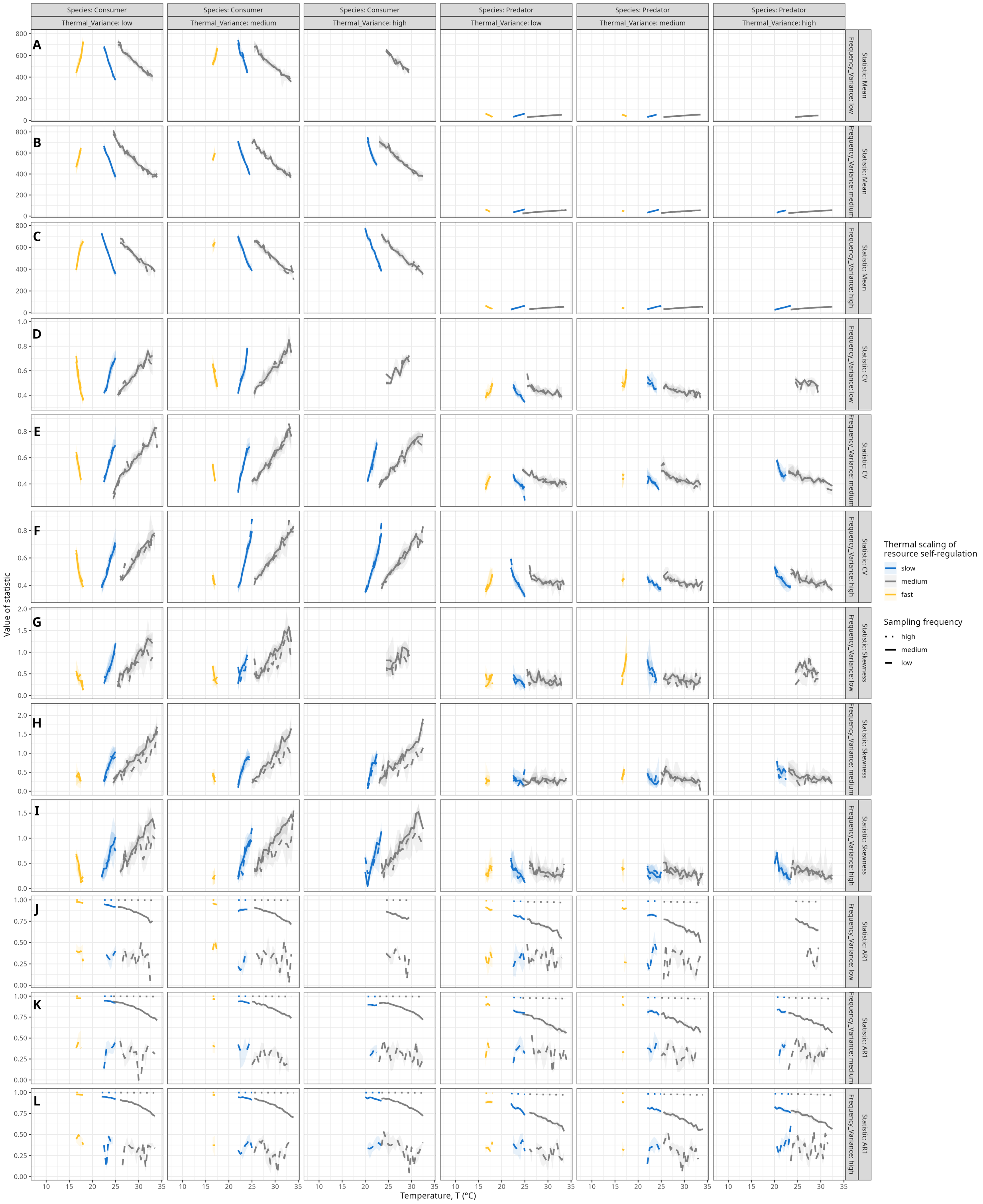


Figure S9: Early warning signals calculated from all stochastic time series in which there was three-species coexistence at $t = 25000$ across species (global x-axis, first level), temperatures (local x-axes), standard deviation of the thermal white noise $\sigma$ (global x-axis, second level; low: 0.5°C, medium: 2°C, high: 5°C), frequency of the thermal white noise $w$ (global y-axis, second level; low: 100, medium: 10, high: 1 time steps), activation energies of resource self-regulation $E_\alpha$ (colors, values as above), and subsampling frequencies (linetype; high: $s = 10$, medium: $s = 100$, low: $s = 1000$ time steps). Lines and shaded regions represent the median and interquartile range across 100 replicate runs of the stochastic simulations. Skewness and AR1 are expressed in absolute values. Initial population densities were set at the scenario-specific deterministic equilibrium density. Other parameter settings are: $r(T_0) = 0.1, \alpha(T_0) = 0.00001, \beta = 3, \gamma = 0.5, \delta = 10, e = 0.05, d(T_0) = 0.01$.
